# Population size and the length of the chromosome blocks identical by descent over generations

**DOI:** 10.1101/032482

**Authors:** Mathieu Tiret, Frédéric Hospital

## Abstract

In all populations, as the time runs, crossovers break apart ancestor haplotypes, forming smaller blocks at each generation. Some blocks, and eventually all of them, become identical by descent because of the genetic drift. We have in this paper developed and benchmarked a theoretical prediction of the mean length of such blocks and used it to study a simple population model assuming panmixia, no selfing and drift as the only evolutionary pressure. Besides, we have on the one hand derived, for any user defined error threshold, the range of the parameters this prediction is reliable for, and on the other hand shown that the mean length remains constant over time in ideally large populations.

---

Identity by descent (IBD) was formally defined for two alleles only, and the definition was subsequently extended to a pair of chromosome segments. Two segments are said to be identical by descent if they are copies of a common ancestor segment without having undergone any mutation. Eventually, Stam (1980) proposed to widen the analyses of identity by descent into an entire population by studying every individual as a pair of chromosome segments. Henceforth, a portion of the chromosome will be referenced as a *segment* and a portion of a segment will be referenced as a *block*.

In a population undergoing genetic drift, as the time runs, crossovers break apart the ancestor haplotypes, forming smaller blocks at each generation. Some blocks might be identical by descent, in which case they are called "IBD blocks". In this paper, we will focus on 1) the distribution in the population of the length of an IBD block, and 2) the distribution of the number and the total length of IBD blocks per individual.

The theoretical framework dealing with these distributions uses a particular population model in order to take account of recombinations: the individuals of this population are diploid, and, following the idea of Stam (1980), they are modeled as a pair of homologous segments with a fixed length *L* (in Morgan). The population size, denoted *N*, is considered constant over generations, there is no evolutionary pressure but the genetic drift, and the generations do not overlap. Moreover, the segments of the founder – or initial – population are all different, that is, none of the 2*N* segments are identical by descent. Finally, the model assumes panmixia without selfing as in Stam (1980), so that two homologous segments of an individual are necessarily derived from two different individuals.

In this model, a segment can be considered as either a discrete or a continuous object. These two ways of modeling are rarely equivalent (Bickeböller and Thompson 1996a,b), and it is not always obvious to know which model should be used. To make the mathematical analyses easier, the segment is here considered as a continuous object: the recombination process can thus be modeled with a classic Poisson process with a rate equal to 1, neglecting crossover interference (Fisher 1949, 1954; Stam 1980; Donnelly 1983; Bickeböller and Thompson 1996a,b; Ball and Stefanov 2005; Chapman and Thompson 2002, 2003; Cannings 2003; Martin and Hospital 2011).

## Theory of junctions

In order to describe the transmission over time of IBD blocks in such population, Fisher (1949) developed the so-called theory of junctions. A *junction* is here a crossover point delimiting two blocks coming from different founders. When considering two segments, it is possible to distinguish two types of junctions: *external junctions*, which are the edges of IBD blocks, and *internal junctions*, which are the other junctions (see Figure 1).

**Figure 1.**
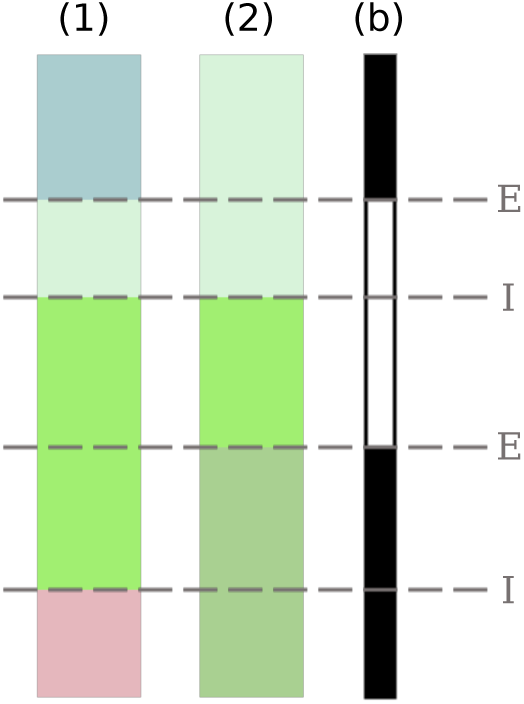
Two segments (1) and (2) of an individual, some time after founding. The different colors represent different founder segments. The white and black bar on the right indicates IBD (white) and non IBD (black) blocks, and each dotted line indicates a junction denoted either E if it is an external junction, or I if it is an internal junction.

An edge of an IBD block is either an external junction or one of the segment edges. Then, knowing the segment length and the number of external junctions, it is possible to infer the distribution in the population of the length of an IBD block. Let *J_N,t_* be the expected number of IBD block edges per individual at generation t. In order to compute *J_N,t_*, Fisher (1949) introduced *H_N,t_* and *Z_N,t_*, which are respectively the expected non IBD proportion of a segment and the expected number of external junctions per Morgan *within the segment* (that is, excluding the segment edges). Besides, let us call IBD segment edge (ISE) a segment edge that is also an edge of an IBD block, and *I_N,t_* the expected number of ISE. Fisher (1949) estimated *I_N,t_* as 2 · (1 − *H_N,t_*) and deduced the following equation:

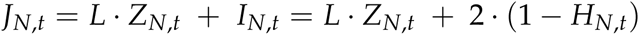

Therefore, the expected number *M_N,t_* of IBD blocks per individual is (Fisher 1949):

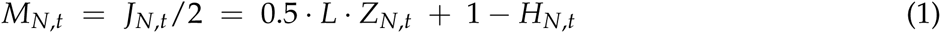

Fisher (1949) figured out a theoretical expression of both *H_N,t_* and *Z_N,t_* only for some very particular cases of relatedness shared by the individuals because of the complexity of the problem. Using identity relations between three genes and their recurrence relations, Stam (1980) derived an expression of *H_N,t_* and *Z_N,t_* for any case of relatedness (see Appendix A for the their formulation). We recall that all these expressions stand for one individual.

Thereby, Fisher (1949) and Stam (1980) described the identity by descent in a population at a *specific time*. Our objective here is to study the evolution of the identity by descent *over time* with a simple population model. To this end, we have developed and benchmarked a prediction of the expected length of an IBD block.

## Model and methods

### Simulations

We have implemented a program generating pseudo-data to compare the predictions with. This program simulates over generations the aforementioned population model. Here, a segment is modeled not as a set of nucleotides but as a continuous object, and hence the program records on the one hand the starting and ending edges of each block and on the other hand the origin of this block, which is the label of the founder segment this block belonged to.

With this program, we have simulated 10000 replicates over 3000 generations, with 10 different population sizes (from 10 to 100 individuals) and with a segment length ranging from 1 to 5 Morgan. The program has been implemented in C++ (version C++11, compiled with g++4.9.2), and graphical outputs have been obtained with R (version 3.2.2).

### Length of identical by descent blocks

Let *X_N,t_* be the expected length of one IBD block. Stam (1980) and Chapman and Thompson (2003), assuming both that the dispatching of junctions over a genome followed a stationary process, obtained the following formulation of *X_N,t_*:

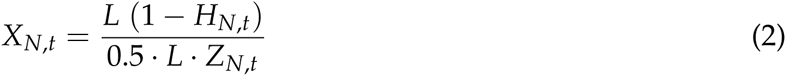

There is however a problem with this assumption: *X_N,t_* as formulated in equation (2) tends to infinity as *t* tends to infinity, whereas the length of an IBD block necessarily ranges from 0 to *L*. The problem comes from *Z_N,t_* that tends to zero as *t* tends to infinity (see Appendix A). The formulation of *Z_N,t_* is correct though, and its asymptotic behavior is indeed expected: genetic drift makes every segment identical after a while, making thus the number of external junctions *Z_N,t_* tend towards zero. The use, however, of the multiplicative inverse of *Z_N,t_* is incorrect, implying that the stationary process assumption is seemingly incompatible with a study of *X_N,t_* over time.

Using equation (1), it is possible to derive another expression of *X_N,t_*:

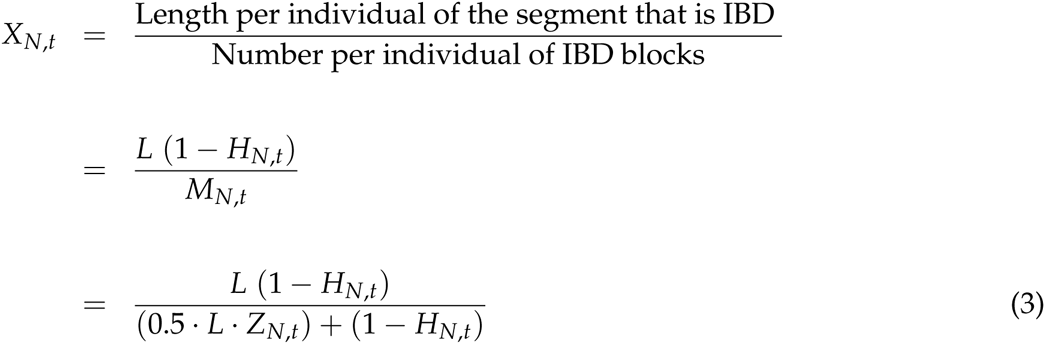

For the problem mentioned above, only equation (3) will hereafter be considered.

### Predictions and observations

The theoretical predictions used in this paper are summarized in Table 1. We recall that, according to equation (3), *X_N,t_* is a combination of *H_N,t_*, *Z_N,t_* and *I_N,t_*.

**Table 1.** Predictions and observations.

| Description | prediction | observation |
| --- | --- | --- |
| Non IBD proportion | $\hat{H}_{N,t}$ | $h_{N,t}$ |
| Number of external junctions | $\hat{Z}_{N,t}$ | $z_{N,t}$ |
| Number of ISE | $\hat{I}_{N,t}$ | $i_{N,t}$ |
| Length of one IBD block | $\hat{X}_{N,t}$ | $x_{N,t}$ |

Of all observations, only the average length *x_N,t_* of an IBD block could be defined in different ways. We have chosen here to define *x_N,t_* as:

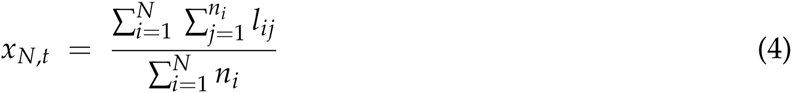

with *n_i_* the number of IBD blocks in the i^th^ individual and *l_ij_* the length of the j^th^ block in the i individual. Thus, *x_N,t_* is defined not as an individual wise but as a population wise observation.

### The focused range of time and the prediction error

We can see on Figure 2 and 3 that *X_N,t_* begins with a high peak, due to the fact that all the founder segments are different. Since we are not interested in this artifactual peak, we will hereafter focus on a range of time starting at *T_N,min_*, the time at which *x_N,t_* reaches its minimum after the peak, and ending at generation 3000. The latter is a fixed value, and the former mainly depends on the population size (see Appendix B).

**Figure 2.**
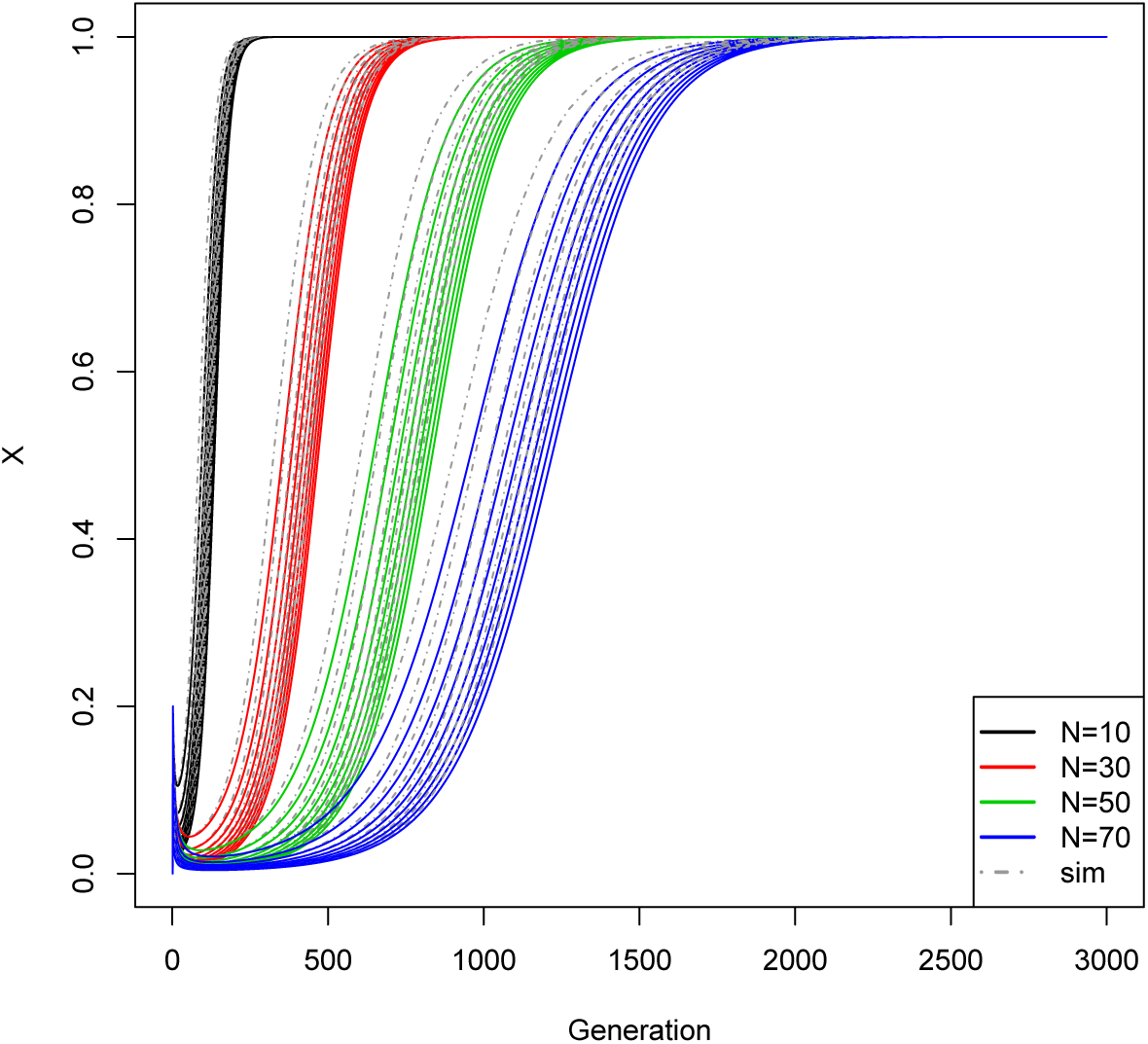
Comparing theoretical prediction and simulation values of *X_N,t_*, with a population size of 10 (black line), 30 (red line), 50 (green line) or 70 individuals (blue line). The solid lines are the estimations, and the dotted lines are the observations (10000 replicates, denoted `sim` in the legend). The different lines of the same color corresponds to the different segment length, ranging from 1 to 5 Morgan.

**Figure 3.**
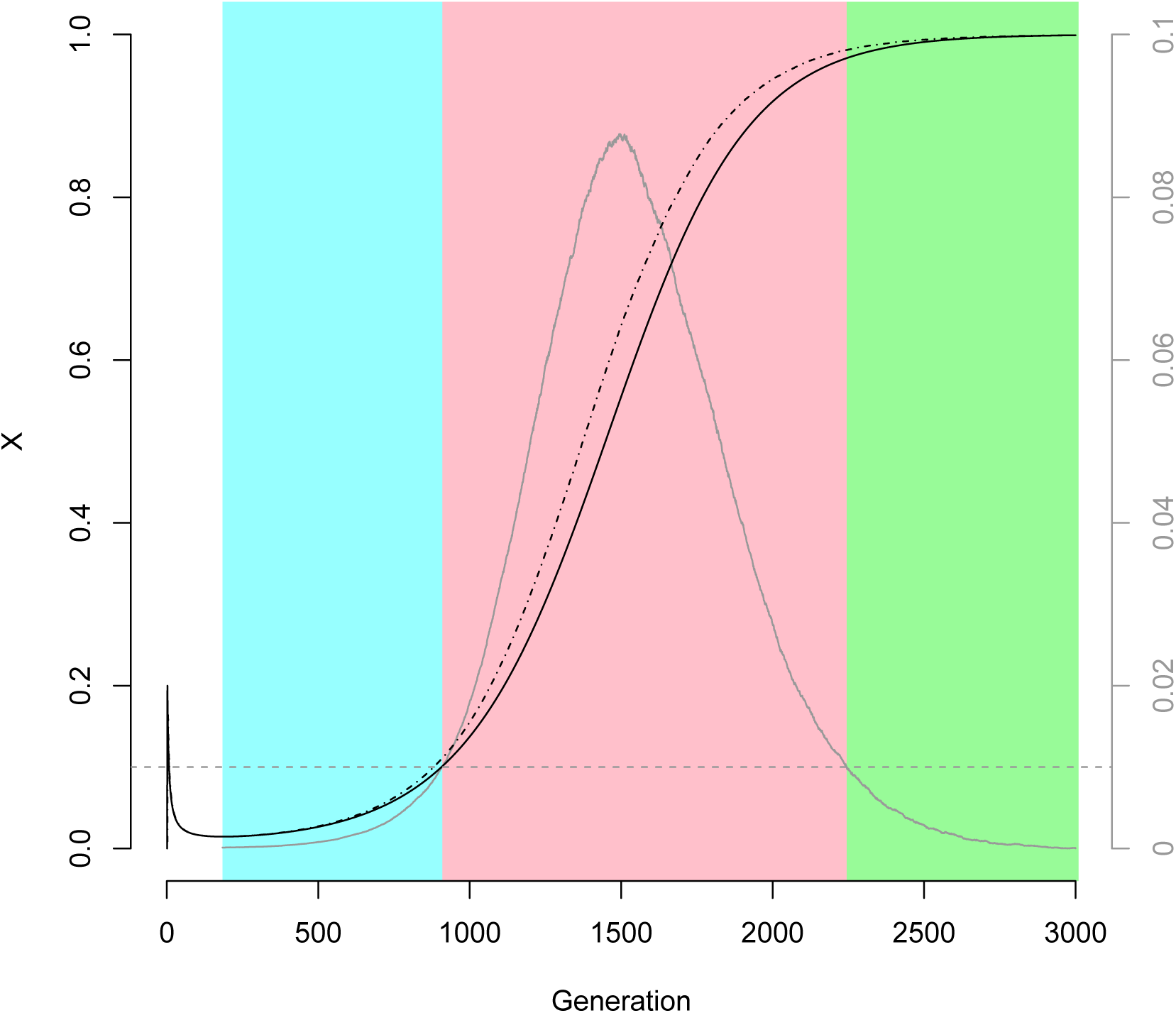
A superposition of the prediction error 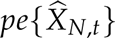, 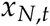 and 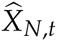 for a population of 100 individuals, *α* of 0.01 (the dashed horizontal line) and a segment length of 1 Morgan. The black solid line is *α*, the black dotted line is *x_N,t_* and the solid gray line is the prediction error 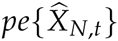, scaled for visual purposes. Its axis is on the right. The blue region corresponds to the first phase, the pink region to the transition phase and the green region to the final phase.

The fit between predictions and observations was ascertained using the prediction error *pe*. With 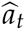 the value at generation *t* of a prediction and *a_t_* the value of the corresponding observation, we have 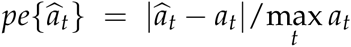.

We have also provided the average value over time of the prediction error, denoted 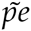. Our main interest being *X_N,t_*, we have searched which component of *X_N,t_* had the greatest prediction error: *H_N,t_, Z_N,t_, I_N,t_* or their combination.

## Results

### The prediction error

Figure 2 shows the prediction and the observation of *X_N,t_* over time, for various population sizes and segment lengths. The average prediction error over *T_N,min_* to generation 3000 are summarized in Table 2.

**Table 2.** The thousandfold value of the average prediction error 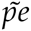 of the different predictions *H_N,t_*, *Z_N,t_*, *I_N,t_* and *X_N,t_*.

| $N$ | 10 | 20 | 30 | 40 | 50 | 60 | 70 | 80 | 90 | 100 |
| --- | --- | --- | --- | --- | --- | --- | --- | --- | --- | --- |
| $\hat{H}_{N,t}$ | 5.96 | 2.90 | 1.92 | 1.26 | 1.01 | 0.93 | 0.75 | 0.64 | 0.55 | 0.48 |
| $\hat{Z}_{N,t}$ | 6.73 | 3.20 | 2.41 | 1.40 | 1.13 | 1.17 | 0.94 | 0.78 | 0.70 | 0.59 |
| $\hat{I}_{N,t}$ | 2.81 | 1.09 | 0.82 | 0.60 | 0.55 | 0.46 | 0.47 | 0.53 | 0.41 | 0.48 |
| $\hat{X}_{N,t}$ | 48.11 | 41.12 | 35.88 | 34.54 | 32.92 | 30.96 | 29.58 | 28.82 | 28.42 | 27.67 |

Table 2 shows that the average prediction error 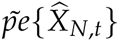 is at least ten times greater than the other average prediction errors, whatever the population size: it seems that a large part of 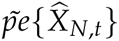 comes from the combination of *H_N,t_*, *Z_N,t_* and *I_N,t_* rather than from each prediction.

We have plotted the prediction error 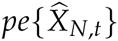 on Figure 3. This figure shows that the error is not constant over time, and has rather a pattern divisible in three phases. Let *α* be a fixed error threshold, ranging from 0 to 1. We define the *first phase* as the phase during which 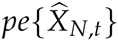 is less than *α*; then the *transition phase* during which 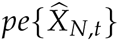 is greater than a; finally, the *final phase* during which 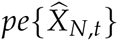 is less than *α* again. Figure 3 shows that the transition phase, during which 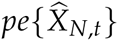 is the greatest, corresponds to the increasing phase of *X_N,t_*.

Figure 3 also shows that 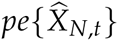 is almost a Gauss-like function and so a normal density function, up to a multiplicative constant. After having fitted 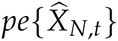 for each population size to a normal gaussian function (see Appendix C), we have deduced the parameters of this function and summarized them in Table 3.

**Table 3.** The parameters (the mean *μ* and the standard deviation *σ*) of the fitted normal distribution and the R-squared (in percent) of this fitting.

| N | 10 | 20 | 30 | 40 | 50 | 60 | 70 | 80 | 90 | 100 |
| --- | --- | --- | --- | --- | --- | --- | --- | --- | --- | --- |
| $\mu$ | 124.67 | 277.28 | 442.39 | 614.83 | 795.89 | 979.67 | 1167.78 | 1357.06 | 1554.44 | 1749.39 |
| $\sigma$ | 29.67 | 58.83 | 86.28 | 112.39 | 142.44 | 168.11 | 193.78 | 220.50 | 251.00 | 277.61 |
| $R^2$ | 98.88 | 98.97 | 98.86 | 98.54 | 98.76 | 98.45 | 98.17 | 98.21 | 98.40 | 98.13 |

According to this fitting and for any value of *α*, we have finally derived the beginning time *T*_1_*_,α_* and the ending time *T*_2_*_,α_* of the transition phase (summarized in Table 4). Between *T_N,min_* and *T*_1_*_,α_* and from *T*_2_*_,α_* until generation 3000, the average prediction error is less than *α*. Furthermore, using the linear regressions of *μ* and *σ*, we have derived that the 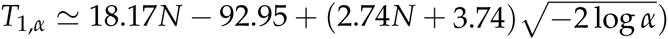 (for further details, see Appendix C):

**Table 4.**
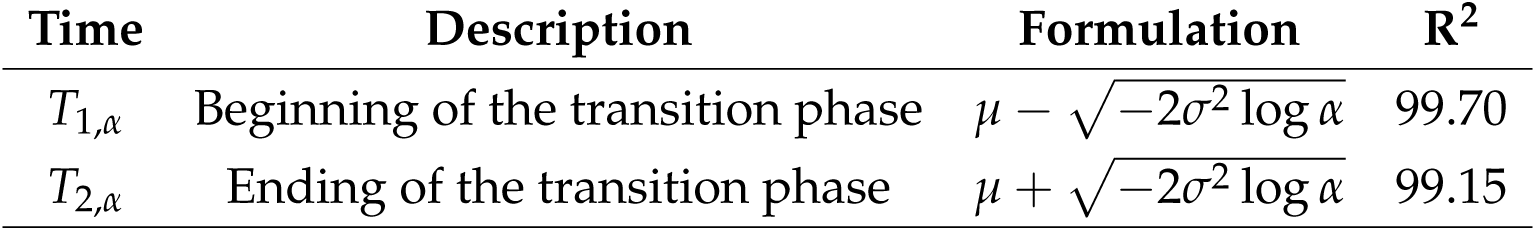
The beginning and the ending times of the transition phase, their formulation and the R-squared of the formulation. *α* is a real number ranging from 0 to 1, *μ* and *α* are respectively the mean and the standard deviation of the fitted normal distribution.

### The minimum length of IBD blocks

Figure 3 shows that our prediction of the expected length of an IBD block is reliable during the first phase, between *T_N,min_* and *T*_1,_*_α_*. During this phase, *X_N,t_* increases only slightly, hence its narrow value range. Besides, the first phase lasts longer as the population size increases, since *T*_1,_*_α_* increases ten times faster than *T_N,min_* does (see Appendix B). Consequently, for a fixed threshold α, the larger a population is, the flatter *X_N,t_* seems to be. The minimum value of *X_N,t_* during the first phase can be estimated as 3/2*N* according to the method of Newton (see Appendix B).

## Discussion

Knowing that the minimum value of *X_N,t_* during the first phase can be estimated as 3/2*N*, *X_N,t_* could therefore be, in an ideally large population, considered constant and equal to 3/2*N* during the first phase. As Table 3 and Table 4 show, the duration of the first phase rapidly increases with the population size, flattening *X_N,t_* and making the ideally large population assumption stand even for a population with 100 individuals, whose first phase lasts about 1,600 generations. It is hence consistent to assume that most populations deriving from highly diverse founders are likely to be in the first phase nowadays, only if it is consistent to assume the model of Haldane, a constant population size and that genetic drift is the only evolutionary pressure.

Mutations were neglected here because the genome was modeled as a continuous object. Indeed, mutations are points, and punctual items do not exist in such continuous models. Compared with the discrete approach, the continuous approach has the advantage of easing the mathematical analyses, but in counterpart it has the shortcoming to assume that there is at the first sight no mutation (it is possible to extend the continuous approach with the infinite allele or the infinite site assumption though), whereas the occurrence of recombination is on average of the same order of magnitude as the occurrence of mutations (the recombination rate is around 10^−8^ per nucleotide per generation for humans). Neglecting mutations is therefore an important limitation to overcome in the future.

Finally, an extension of this framework will consist on the one hand in theoretically determining the variance of the distribution of *X_N,t_* and on the other hand in focusing on the impact of the founder population and its structure: assuming that every founder segment is different, as we did here (according to Stam 1980), is more than unlikely in a real population. It will be important in further studies to assess whether this structure changes the whole dynamic or only, as for a Markovian process, the beginning of the process.

## Acknowledgements

This work has been supported by grants from the metaprogram SelGen, Institut Nationale de Recherche Agronomique, INRA (to F.H.). We are also grateful to the genotoul bioinformatics platform Toulouse Midi-Pyrenees (Bioinfo Genotoul) for providing help and computing resources.

## Appendices

### A. Mathematical formulation of *H* and *Z*

The exact expression of *H_N,t_* and *Z_N,t_* was described by Stam (1980) and reads:

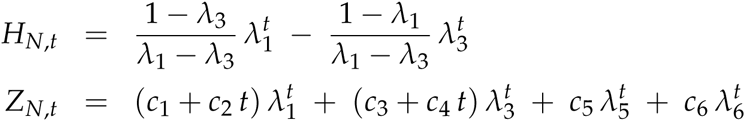

with *c_i_*’s and *λ_i_*’s values depending on *N*, as follows:

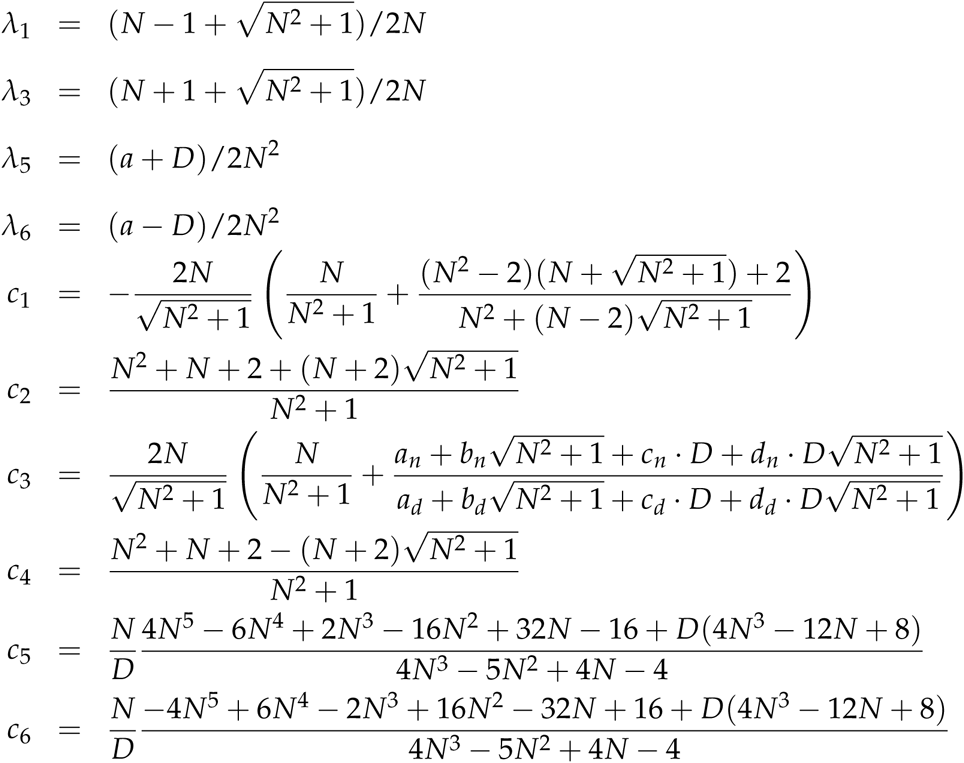

with

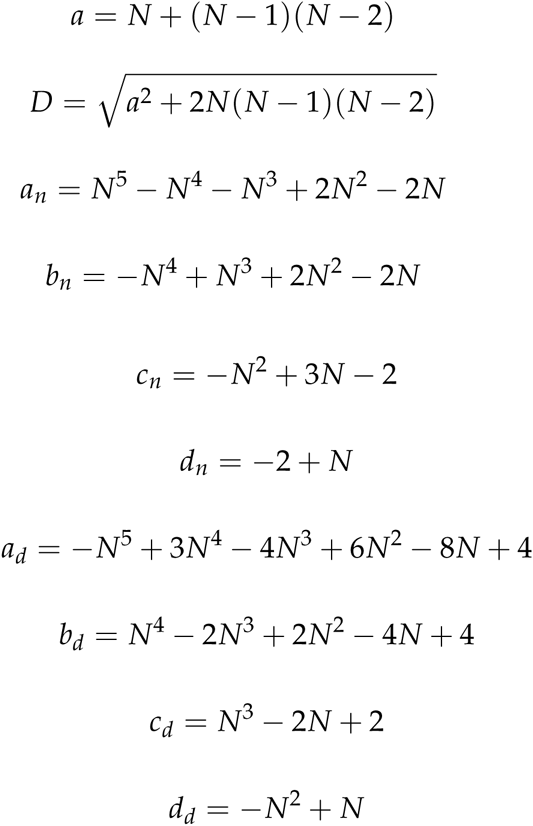

To ease mathematical analyses, we have also derived the series expansion as *N* tends towards infinity of those coefficients:

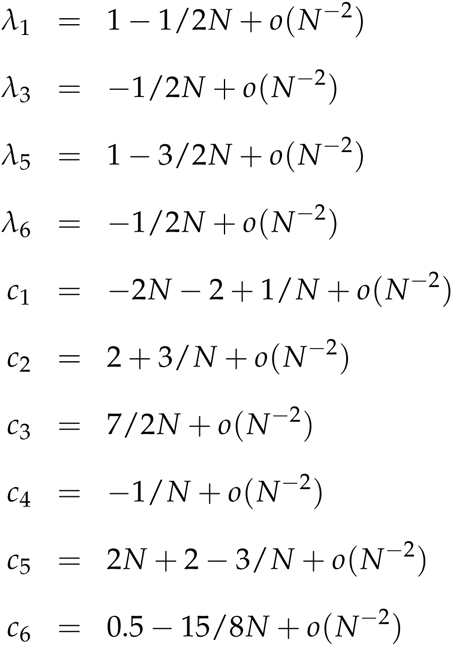

Hence, we can derive that:

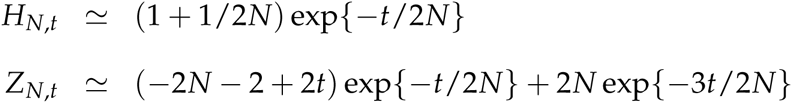

It is now trivial to deduce that *Z_N,t_* tends towards zero as t tends towards infinity.

### B. The minimum value of *X*

We have derived with a linear regression on simulations that *T_N,min_* ≃ 1.87*N* − 1.27 with an adjusted R-squared of 0.9992 and that 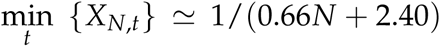 with an adjusted R-squared of 1. Besides, using the Newton's method on theoretical expressions of the predictions, we have derived that 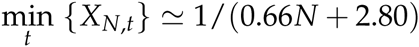. Both approximations of 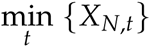 are equivalent and could roughly be approximated as 3/2*N*.

### C. Normal adjustment

We estimated here the parameters *μ* and *σ* of the fitted normal distribution of *pe* as follows:

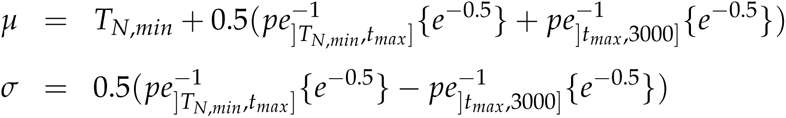

with *t_max_* the time at which *pe* reaches its maximum and 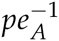 the inverse function of *pe* on *A*. The coefficient of determination *R*^2^ of the normal fitting was computed as follows:

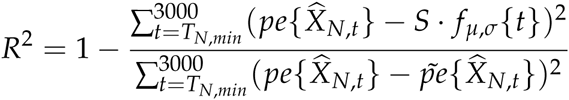

with *S* a corrective coefficient (equal to 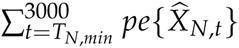) to make *pe* have its integration over the real numbers equal to 1, *f_μ,σ_* the density function of the normal distribution with a mean *μ* and a standard deviation *σ*, and 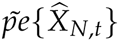 the average value of *pe* over *T_N,min_* to the generation 3000.

With a linear regression, we have derived that *μ* ≃ 18.17*N* − 92.95 with an adjusted R-squared of 0.9773 and *σ* ≃ 2.74*N* + 3.47 with an adjusted R-squared of 0.9959.

